## Supplementary Figures for "Integrative spatial omics reveals distinct tumor-promoting multicellular niches and immunosuppressive mechanisms in Black American and White American patients with TNBC"

### Slide 1
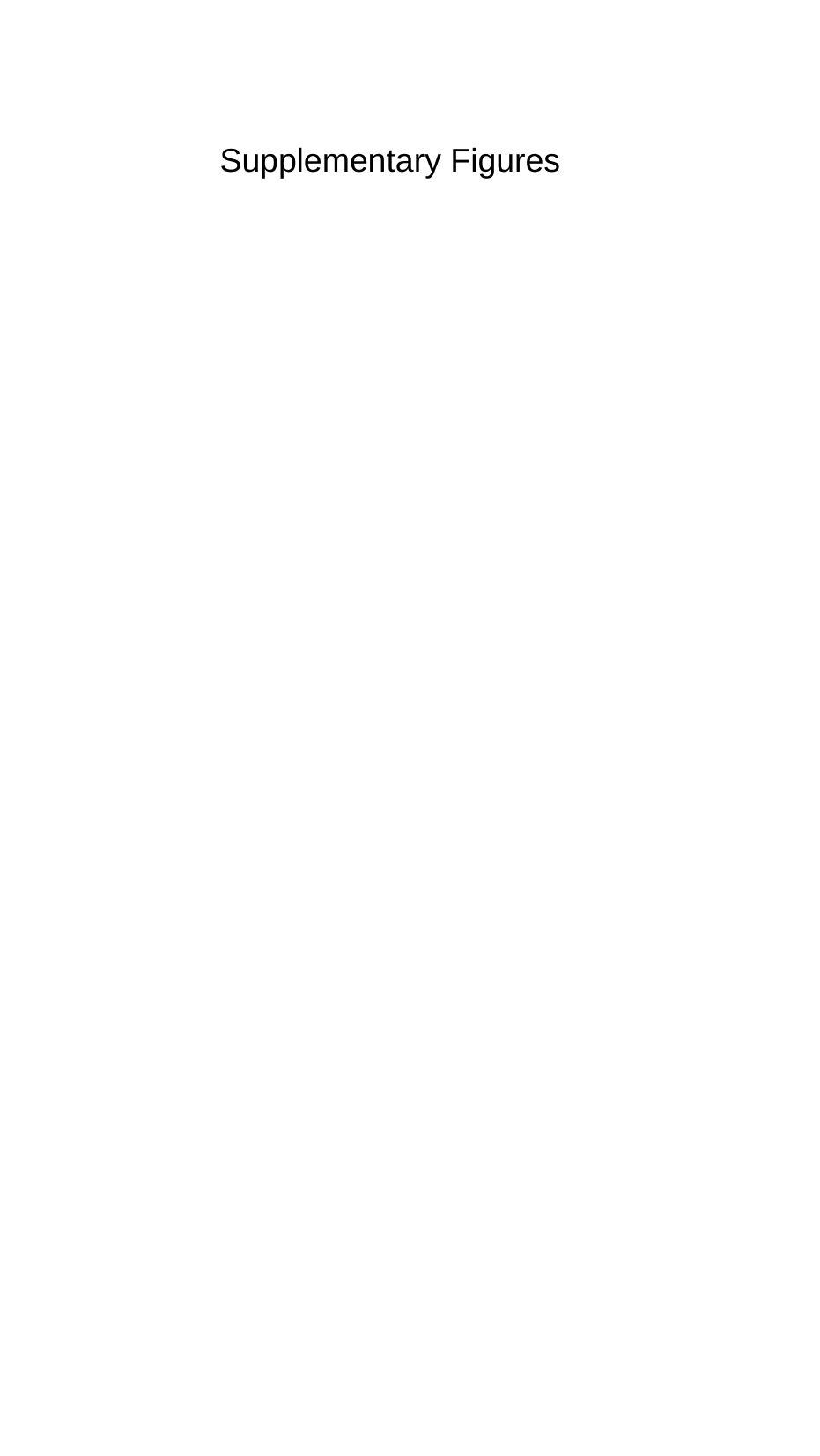

Supplementary Figures

### Slide 2
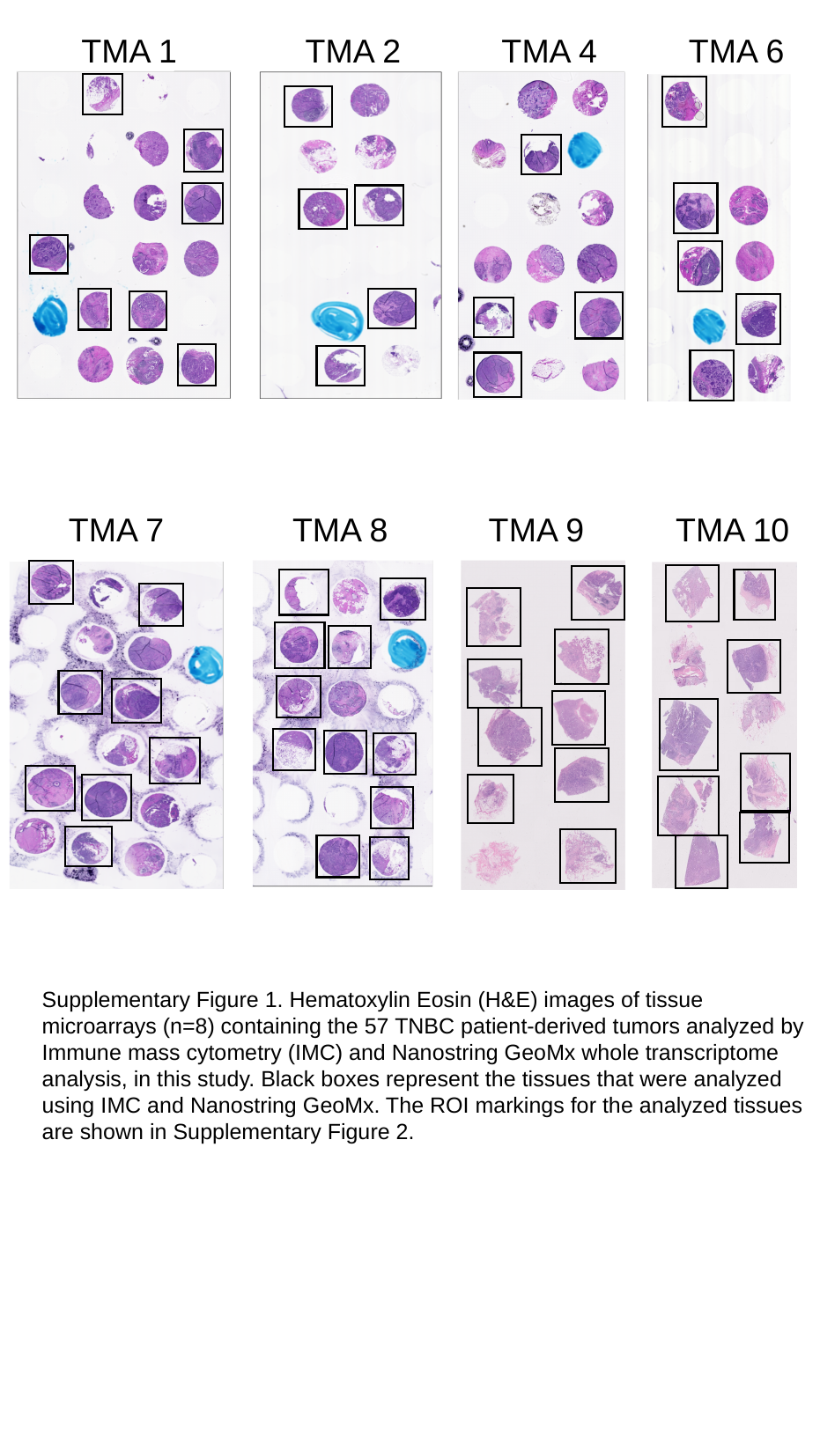

TMA 1 TMA 2 TMA 4 TMA 6
TMA 7 TMA 8 TMA 9 TMA 10
Supplementary Figure 1. Hematoxylin Eosin (H&E) images of tissue microarrays (n=8) containing the 57 TNBC patient-derived tumors analyzed by Immune mass cytometry (IMC) and Nanostring GeoMx whole transcriptome analysis, in this study. Black boxes represent the tissues that were analyzed using IMC and Nanostring GeoMx. The ROI markings for the analyzed tissues are shown in Supplementary Figure 2.

### Slide 3
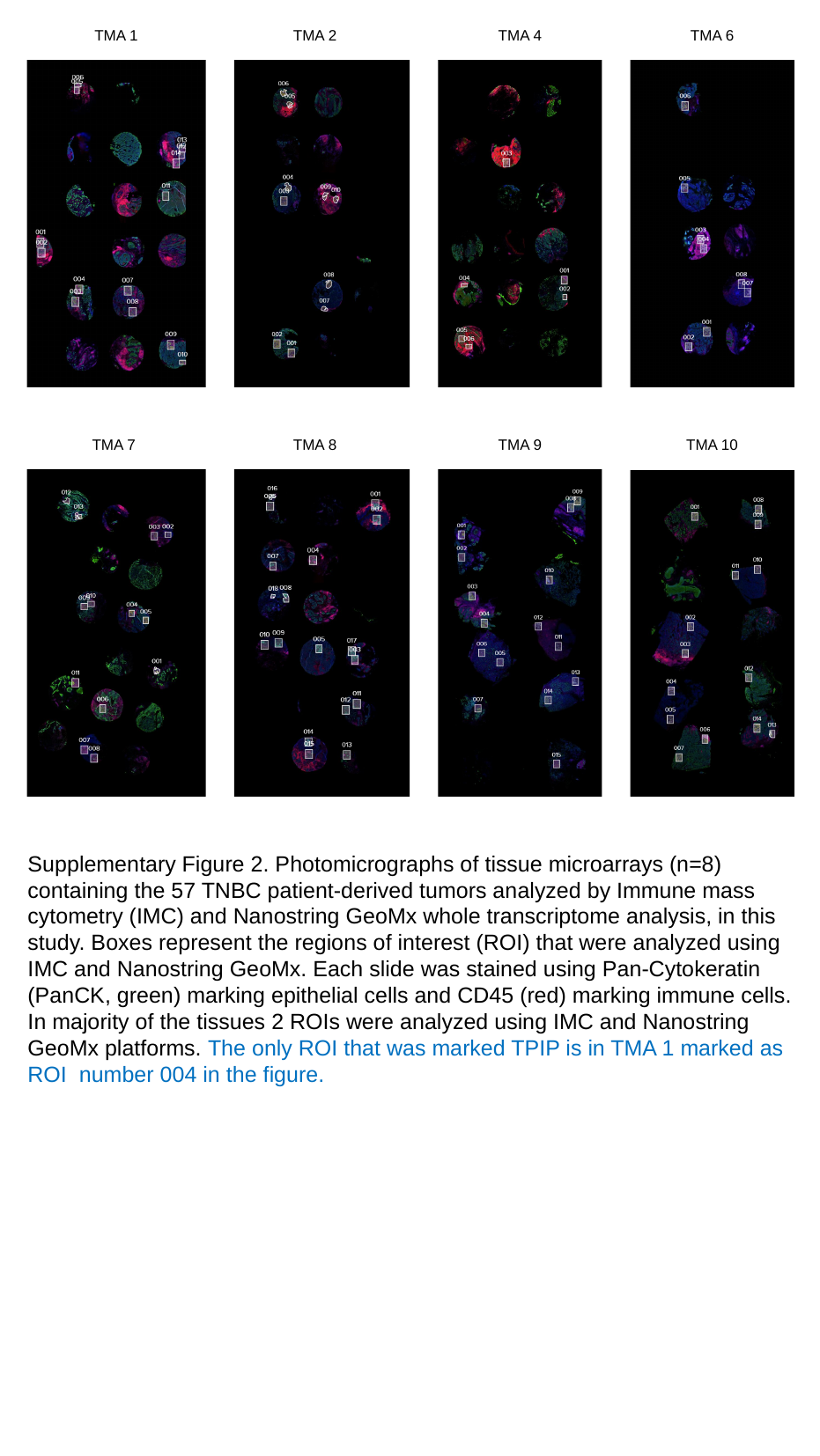

TMA 2
TMA 4
TMA 6
TMA 1
TMA 7
TMA 8
TMA 9
TMA 10
Supplementary Figure 2. Photomicrographs of tissue microarrays (n=8) containing the 57 TNBC patient-derived tumors analyzed by Immune mass cytometry (IMC) and Nanostring GeoMx whole transcriptome analysis, in this study. Boxes represent the regions of interest (ROI) that were analyzed using IMC and Nanostring GeoMx. Each slide was stained using Pan-Cytokeratin (PanCK, green) marking epithelial cells and CD45 (red) marking immune cells. In majority of the tissues 2 ROIs were analyzed using IMC and Nanostring GeoMx platforms. The only ROI that was marked TPIP is in TMA 1 marked as ROI number 004 in the figure.

### Slide 4
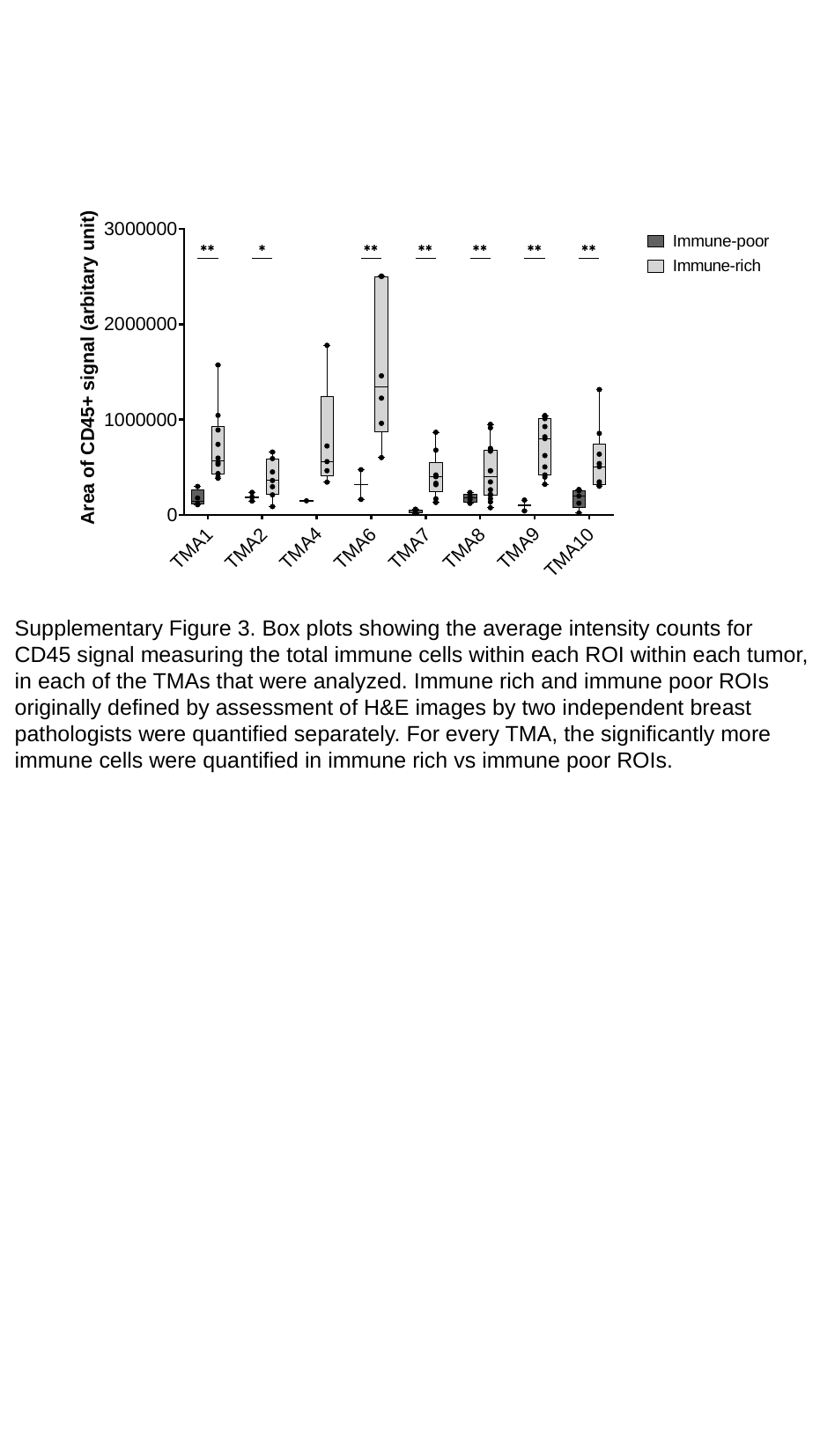

Supplementary Figure 3. Box plots showing the average intensity counts for
CD45 signal measuring the total immune cells within each ROI within each tumor,
in each of the TMAs that were analyzed. Immune rich and immune poor ROIs
originally defined by assessment of H&E images by two independent breast
pathologists were quantified separately. For every TMA, the significantly more
immune cells were quantified in immune rich vs immune poor ROIs.

### Slide 5
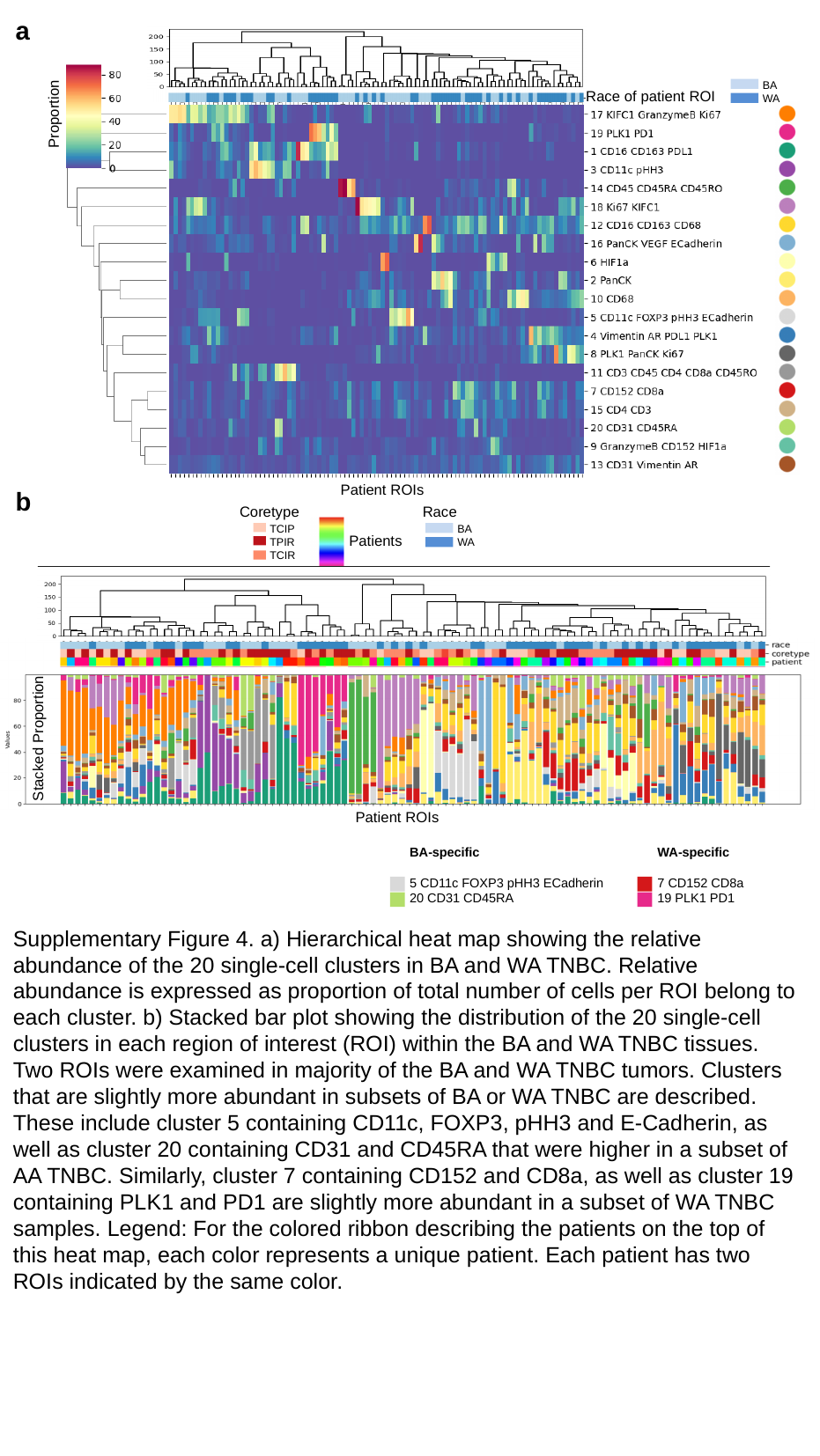

a
BA
WA
Race of patient ROI
Proportion
Patient ROIs
b
Race
Coretype
TCIP
TPIR
TCIR
BA
WA
Patients
Stacked Proportion
Patient ROIs
WA-specific
7 CD152 CD8a
19 PLK1 PD1
BA-specific
5 CD11c FOXP3 pHH3 ECadherin
20 CD31 CD45RA
Supplementary Figure 4. a) Hierarchical heat map showing the relative abundance of the 20 single-cell clusters in BA and WA TNBC. Relative abundance is expressed as proportion of total number of cells per ROI belong to each cluster. b) Stacked bar plot showing the distribution of the 20 single-cell clusters in each region of interest (ROI) within the BA and WA TNBC tissues. Two ROIs were examined in majority of the BA and WA TNBC tumors. Clusters that are slightly more abundant in subsets of BA or WA TNBC are described. These include cluster 5 containing CD11c, FOXP3, pHH3 and E-Cadherin, as well as cluster 20 containing CD31 and CD45RA that were higher in a subset of AA TNBC. Similarly, cluster 7 containing CD152 and CD8a, as well as cluster 19 containing PLK1 and PD1 are slightly more abundant in a subset of WA TNBC samples. Legend: For the colored ribbon describing the patients on the top of this heat map, each color represents a unique patient. Each patient has two ROIs indicated by the same color.

### Slide 6
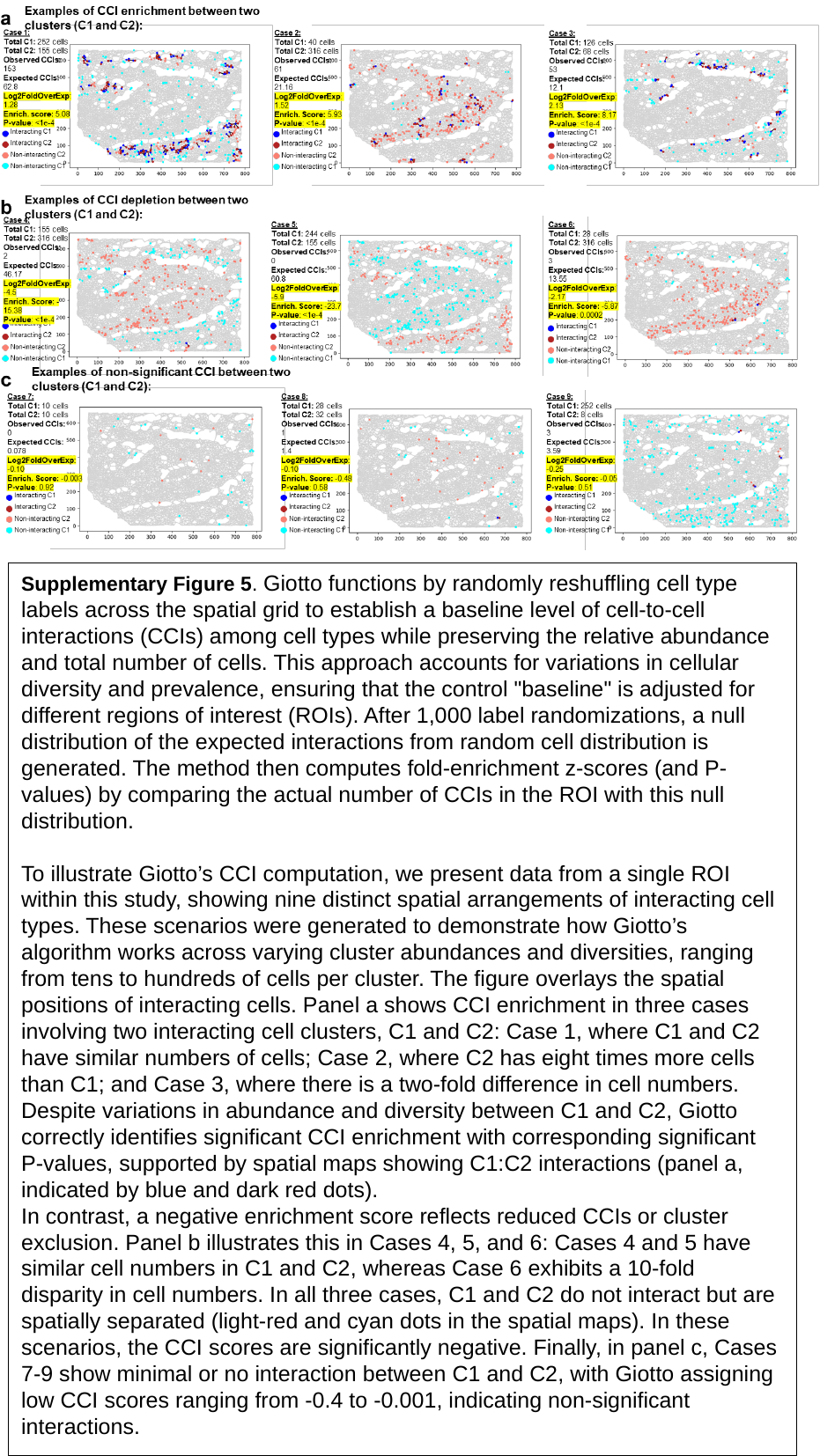

Supplementary Figure 5. Giotto functions by randomly reshuffling cell type labels across the spatial grid to establish a baseline level of cell-to-cell interactions (CCIs) among cell types while preserving the relative abundance and total number of cells. This approach accounts for variations in cellular diversity and prevalence, ensuring that the control "baseline" is adjusted for different regions of interest (ROIs). After 1,000 label randomizations, a null distribution of the expected interactions from random cell distribution is generated. The method then computes fold-enrichment z-scores (and P-values) by comparing the actual number of CCIs in the ROI with this null distribution.
To illustrate Giotto’s CCI computation, we present data from a single ROI within this study, showing nine distinct spatial arrangements of interacting cell types. These scenarios were generated to demonstrate how Giotto’s algorithm works across varying cluster abundances and diversities, ranging from tens to hundreds of cells per cluster. The figure overlays the spatial positions of interacting cells. Panel a shows CCI enrichment in three cases involving two interacting cell clusters, C1 and C2: Case 1, where C1 and C2 have similar numbers of cells; Case 2, where C2 has eight times more cells than C1; and Case 3, where there is a two-fold difference in cell numbers. Despite variations in abundance and diversity between C1 and C2, Giotto correctly identifies significant CCI enrichment with corresponding significant P-values, supported by spatial maps showing C1:C2 interactions (panel a, indicated by blue and dark red dots).
In contrast, a negative enrichment score reflects reduced CCIs or cluster exclusion. Panel b illustrates this in Cases 4, 5, and 6: Cases 4 and 5 have similar cell numbers in C1 and C2, whereas Case 6 exhibits a 10-fold disparity in cell numbers. In all three cases, C1 and C2 do not interact but are spatially separated (light-red and cyan dots in the spatial maps). In these scenarios, the CCI scores are significantly negative. Finally, in panel c, Cases 7-9 show minimal or no interaction between C1 and C2, with Giotto assigning low CCI scores ranging from -0.4 to -0.001, indicating non-significant interactions.

### Slide 7
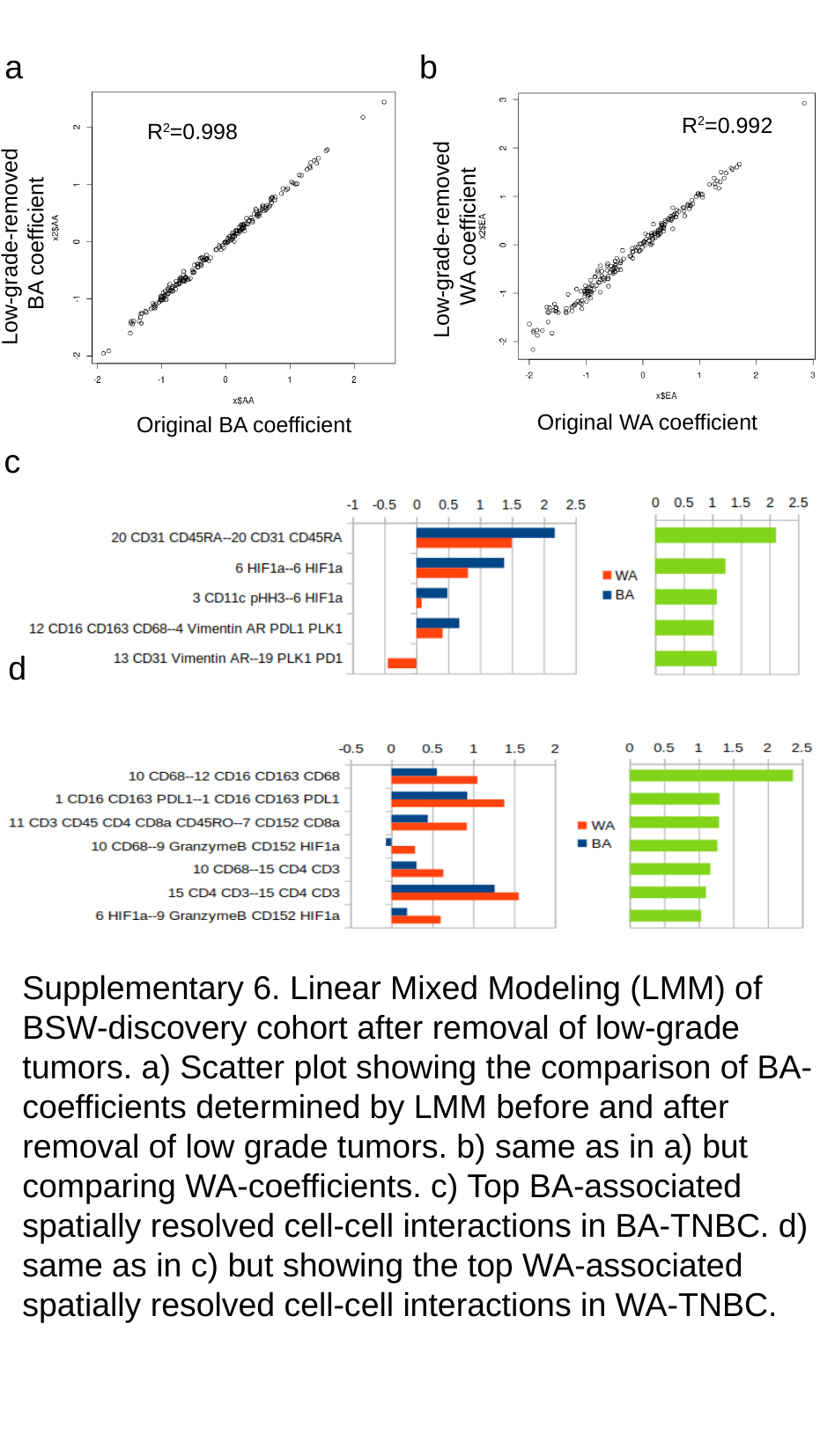

a b
R^2^=0.992
Low-grade-removed
WA coefficient
Original WA coefficient
R^2^=0.998
Low-grade-removed
BA coefficient
Original BA coefficient
c
d
Supplementary 6. Linear Mixed Modeling (LMM) of
BSW-discovery cohort after removal of low-grade
tumors. a) Scatter plot showing the comparison of BA-
coefficients determined by LMM before and after
removal of low grade tumors. b) same as in a) but
comparing WA-coefficients. c) Top BA-associated
spatially resolved cell-cell interactions in BA-TNBC. d)
same as in c) but showing the top WA-associated
spatially resolved cell-cell interactions in WA-TNBC.

### Slide 8
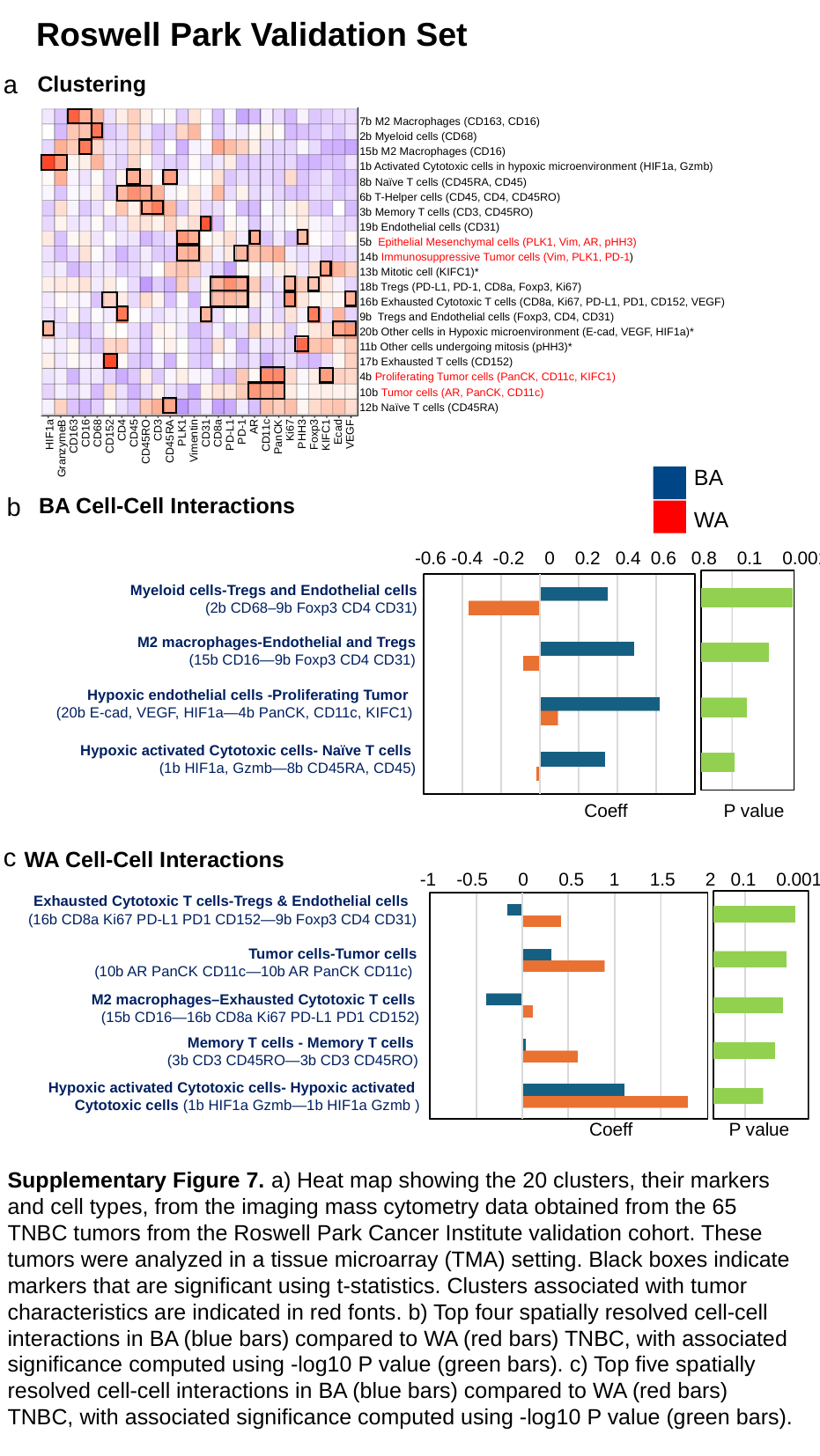

Roswell Park Validation Set
Clustering
a
7b M2 Macrophages (CD163, CD16)
2b Myeloid cells (CD68)
15b M2 Macrophages (CD16)
1b Activated Cytotoxic cells in hypoxic microenvironment (HIF1a, Gzmb)
8b Naïve T cells (CD45RA, CD45)
6b T-Helper cells (CD45, CD4, CD45RO)
3b Memory T cells (CD3, CD45RO)
19b Endothelial cells (CD31)
5b Epithelial Mesenchymal cells (PLK1, Vim, AR, pHH3)
14b Immunosuppressive Tumor cells (Vim, PLK1, PD-1)
13b Mitotic cell (KIFC1)*
18b Tregs (PD-L1, PD-1, CD8a, Foxp3, Ki67)
16b Exhausted Cytotoxic T cells (CD8a, Ki67, PD-L1, PD1, CD152, VEGF)
9b Tregs and Endothelial cells (Foxp3, CD4, CD31)
20b Other cells in Hypoxic microenvironment (E-cad, VEGF, HIF1a)*
11b Other cells undergoing mitosis (pHH3)*
17b Exhausted T cells (CD152)
4b Proliferating Tumor cells (PanCK, CD11c, KIFC1)
10b Tumor cells (AR, PanCK, CD11c)
12b Naïve T cells (CD45RA)
HIF1a
GranzymeB
CD163
CD16
CD68
CD152
CD4
CD45
CD45RO
CD3
CD45RA
PLK1
Vimentin
CD31
CD8a
PD-L1
PD-1
AR
CD11c
PanCK
Ki67
PHH3
Foxp3
KIFC1
Ecad
VEGF
BA
WA
BA Cell-Cell Interactions
b
-0.6 -0.4 -0.2 0 0.2 0.4 0.6 0.8 0.1 0.001
 Myeloid cells-Tregs and Endothelial cells
 (2b CD68–9b Foxp3 CD4 CD31)
 M2 macrophages-Endothelial and Tregs
 (15b CD16—9b Foxp3 CD4 CD31)
 Hypoxic endothelial cells -Proliferating Tumor
 (20b E-cad, VEGF, HIF1a—4b PanCK, CD11c, KIFC1)
 Hypoxic activated Cytotoxic cells- Naïve T cells
 (1b HIF1a, Gzmb—8b CD45RA, CD45)
Coeff P value
WA Cell-Cell Interactions
c
-1 -0.5 0 0.5 1 1.5 2 0.1 0.001
 Exhausted Cytotoxic T cells-Tregs & Endothelial cells
 (16b CD8a Ki67 PD-L1 PD1 CD152—9b Foxp3 CD4 CD31)
 Tumor cells-Tumor cells
 (10b AR PanCK CD11c—10b AR PanCK CD11c)
M2 macrophages–Exhausted Cytotoxic T cells
 (15b CD16—16b CD8a Ki67 PD-L1 PD1 CD152)
 Memory T cells - Memory T cells
 (3b CD3 CD45RO—3b CD3 CD45RO)
 Hypoxic activated Cytotoxic cells- Hypoxic activated
Cytotoxic cells (1b HIF1a Gzmb—1b HIF1a Gzmb )
Coeff P value
Supplementary Figure 7. a) Heat map showing the 20 clusters, their markers and cell types, from the imaging mass cytometry data obtained from the 65 TNBC tumors from the Roswell Park Cancer Institute validation cohort. These tumors were analyzed in a tissue microarray (TMA) setting. Black boxes indicate markers that are significant using t-statistics. Clusters associated with tumor characteristics are indicated in red fonts. b) Top four spatially resolved cell-cell interactions in BA (blue bars) compared to WA (red bars) TNBC, with associated significance computed using -log10 P value (green bars). c) Top five spatially resolved cell-cell interactions in BA (blue bars) compared to WA (red bars) TNBC, with associated significance computed using -log10 P value (green bars).

### Slide 9
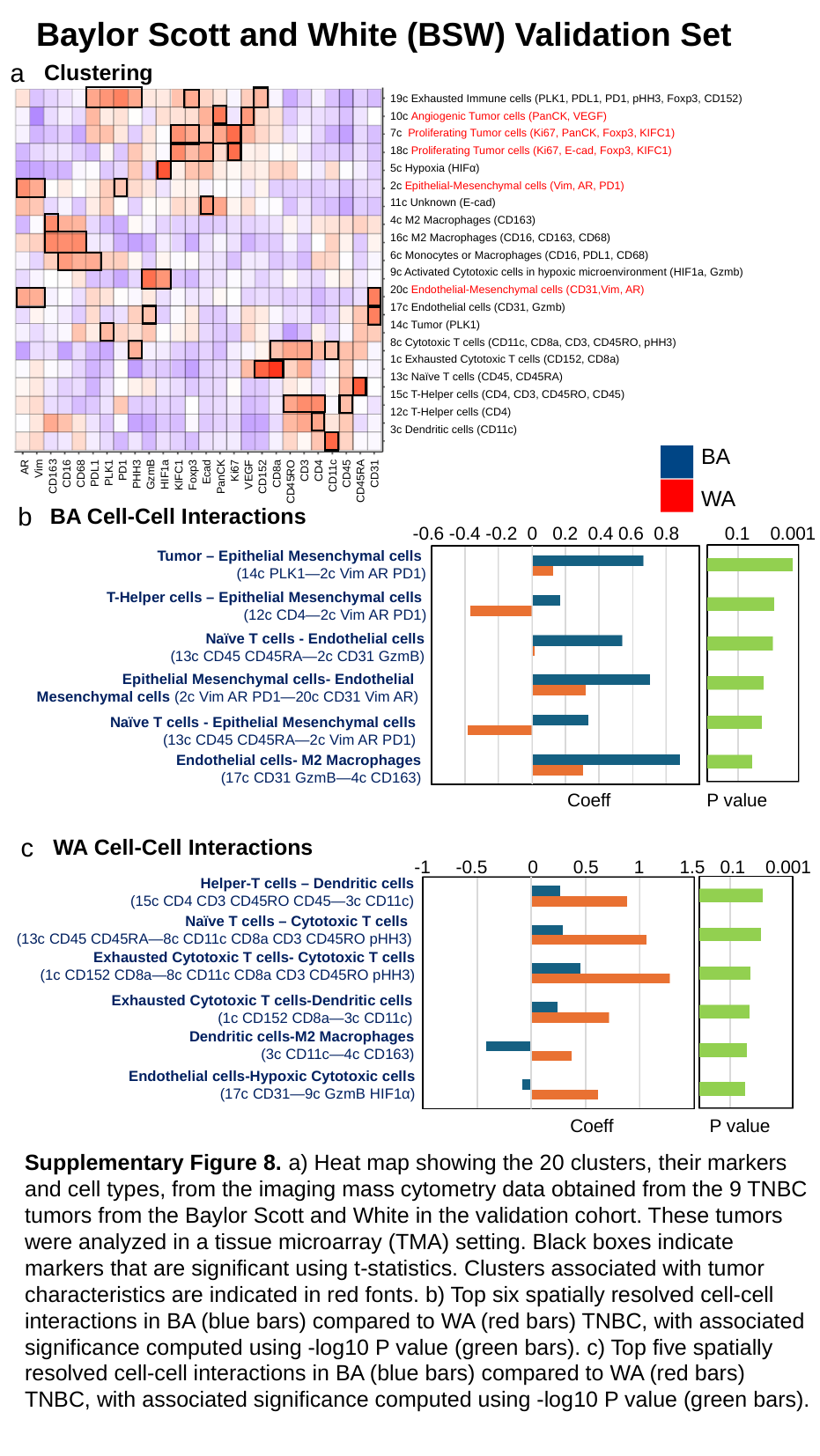

Baylor Scott and White (BSW) Validation Set
Clustering
a
19c Exhausted Immune cells (PLK1, PDL1, PD1, pHH3, Foxp3, CD152)
10c Angiogenic Tumor cells (PanCK, VEGF)
7c Proliferating Tumor cells (Ki67, PanCK, Foxp3, KIFC1)
18c Proliferating Tumor cells (Ki67, E-cad, Foxp3, KIFC1)
5c Hypoxia (HIFα)
2c Epithelial-Mesenchymal cells (Vim, AR, PD1)
11c Unknown (E-cad)
4c M2 Macrophages (CD163)
16c M2 Macrophages (CD16, CD163, CD68)
6c Monocytes or Macrophages (CD16, PDL1, CD68)
9c Activated Cytotoxic cells in hypoxic microenvironment (HIF1a, Gzmb)
20c Endothelial-Mesenchymal cells (CD31,Vim, AR)
17c Endothelial cells (CD31, Gzmb)
14c Tumor (PLK1)
8c Cytotoxic T cells (CD11c, CD8a, CD3, CD45RO, pHH3)
1c Exhausted Cytotoxic T cells (CD152, CD8a)
13c Naïve T cells (CD45, CD45RA)
15c T-Helper cells (CD4, CD3, CD45RO, CD45)
12c T-Helper cells (CD4)
3c Dendritic cells (CD11c)
AR
Vim
CD163
CD16
CD68
PDL1
PLK1
PD1
PHH3
GzmB
HIF1a
KIFC1
Foxp3
Ecad
PanCK
Ki67
VEGF
CD152
CD8a
CD45RO
CD3
CD4
CD11c
CD45
CD45RA
CD31
BA
WA
BA Cell-Cell Interactions
b
-0.6 -0.4 -0.2 0 0.2 0.4 0.6 0.8 0.1 0.001
 Tumor – Epithelial Mesenchymal cells
 (14c PLK1—2c Vim AR PD1)
 T-Helper cells – Epithelial Mesenchymal cells
 (12c CD4—2c Vim AR PD1)
 Naïve T cells - Endothelial cells
 (13c CD45 CD45RA—2c CD31 GzmB)
 Epithelial Mesenchymal cells- Endothelial
Mesenchymal cells (2c Vim AR PD1—20c CD31 Vim AR)
 Naïve T cells - Epithelial Mesenchymal cells
 (13c CD45 CD45RA—2c Vim AR PD1)
Endothelial cells- M2 Macrophages
 (17c CD31 GzmB—4c CD163)
Coeff P value
WA Cell-Cell Interactions
c
-1 -0.5 0 0.5 1 1.5 0.1 0.001
 Helper-T cells – Dendritic cells
 (15c CD4 CD3 CD45RO CD45—3c CD11c)
 Naïve T cells – Cytotoxic T cells
(13c CD45 CD45RA—8c CD11c CD8a CD3 CD45RO pHH3)
 Exhausted Cytotoxic T cells- Cytotoxic T cells
(1c CD152 CD8a—8c CD11c CD8a CD3 CD45RO pHH3)
Exhausted Cytotoxic T cells-Dendritic cells
 (1c CD152 CD8a—3c CD11c)
Dendritic cells-M2 Macrophages
 (3c CD11c—4c CD163)
Endothelial cells-Hypoxic Cytotoxic cells
 (17c CD31—9c GzmB HIF1α)
Coeff P value
Supplementary Figure 8. a) Heat map showing the 20 clusters, their markers and cell types, from the imaging mass cytometry data obtained from the 9 TNBC tumors from the Baylor Scott and White in the validation cohort. These tumors were analyzed in a tissue microarray (TMA) setting. Black boxes indicate markers that are significant using t-statistics. Clusters associated with tumor characteristics are indicated in red fonts. b) Top six spatially resolved cell-cell interactions in BA (blue bars) compared to WA (red bars) TNBC, with associated significance computed using -log10 P value (green bars). c) Top five spatially resolved cell-cell interactions in BA (blue bars) compared to WA (red bars) TNBC, with associated significance computed using -log10 P value (green bars).

### Slide 10
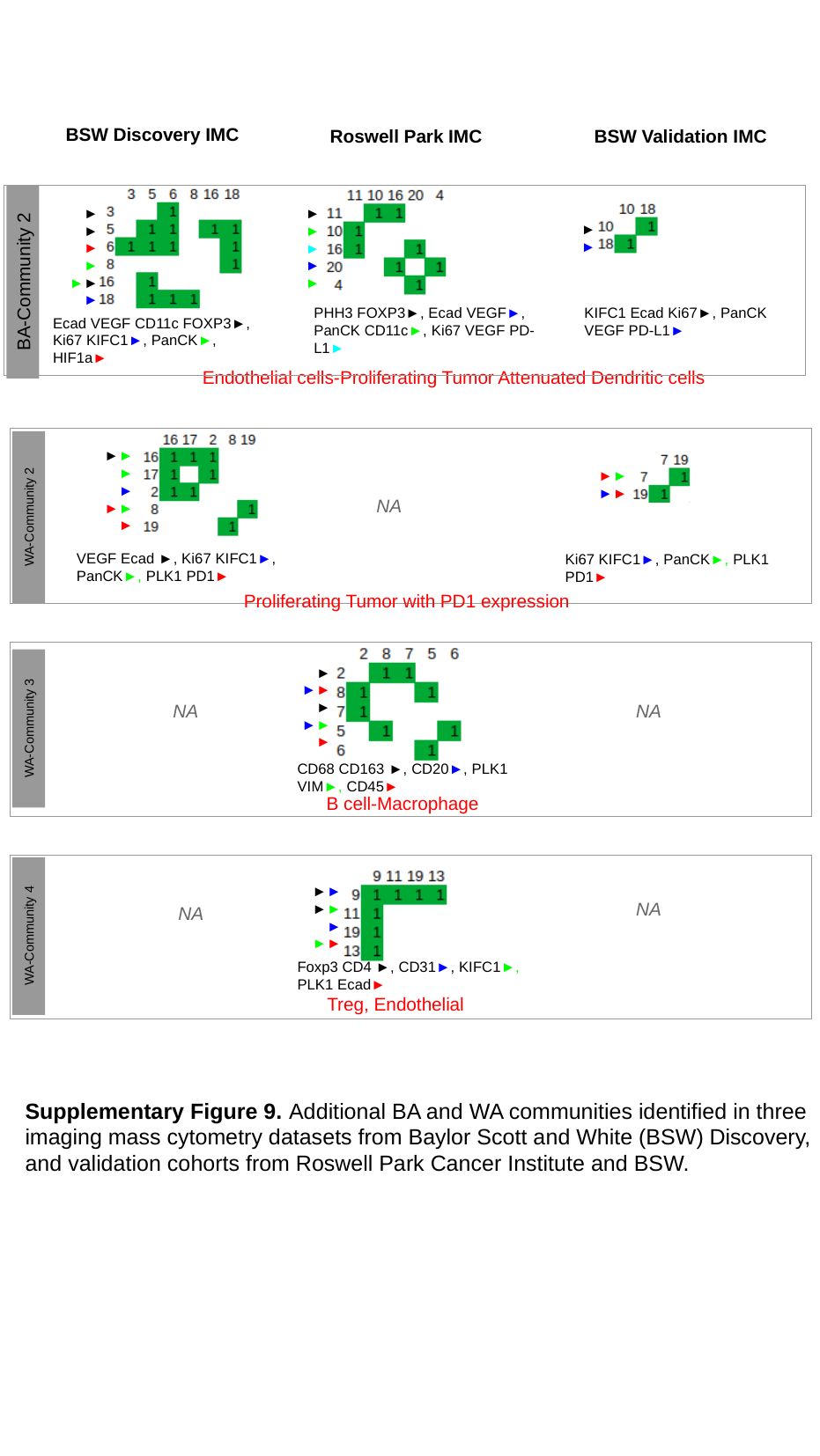

BSW Discovery IMC
Roswell Park IMC
BSW Validation IMC
►
►
►
►
►
►
►
►
►
►
►
►►
►
BA-Community 2
PHH3 FOXP3►, Ecad VEGF►, PanCK CD11c►, Ki67 VEGF PD-L1►
KIFC1 Ecad Ki67►, PanCK VEGF PD-L1►
Ecad VEGF CD11c FOXP3►,
Ki67 KIFC1►, PanCK►, HIF1a►
Endothelial cells-Proliferating Tumor Attenuated Dendritic cells
►►
►
►
►►
►
►►
►►
NA
WA-Community 2
VEGF Ecad ►, Ki67 KIFC1►, PanCK►, PLK1 PD1►
Ki67 KIFC1►, PanCK►, PLK1 PD1►
Proliferating Tumor with PD1 expression
►
►►
►
►►
►
NA
NA
WA-Community 3
CD68 CD163 ►, CD20►, PLK1 VIM►, CD45►
B cell-Macrophage
►►►►►
►►
NA
NA
WA-Community 4
Foxp3 CD4 ►, CD31►, KIFC1►, PLK1 Ecad►
Treg, Endothelial
Supplementary Figure 9. Additional BA and WA communities identified in three
imaging mass cytometry datasets from Baylor Scott and White (BSW) Discovery, and validation cohorts from Roswell Park Cancer Institute and BSW.

### Slide 11
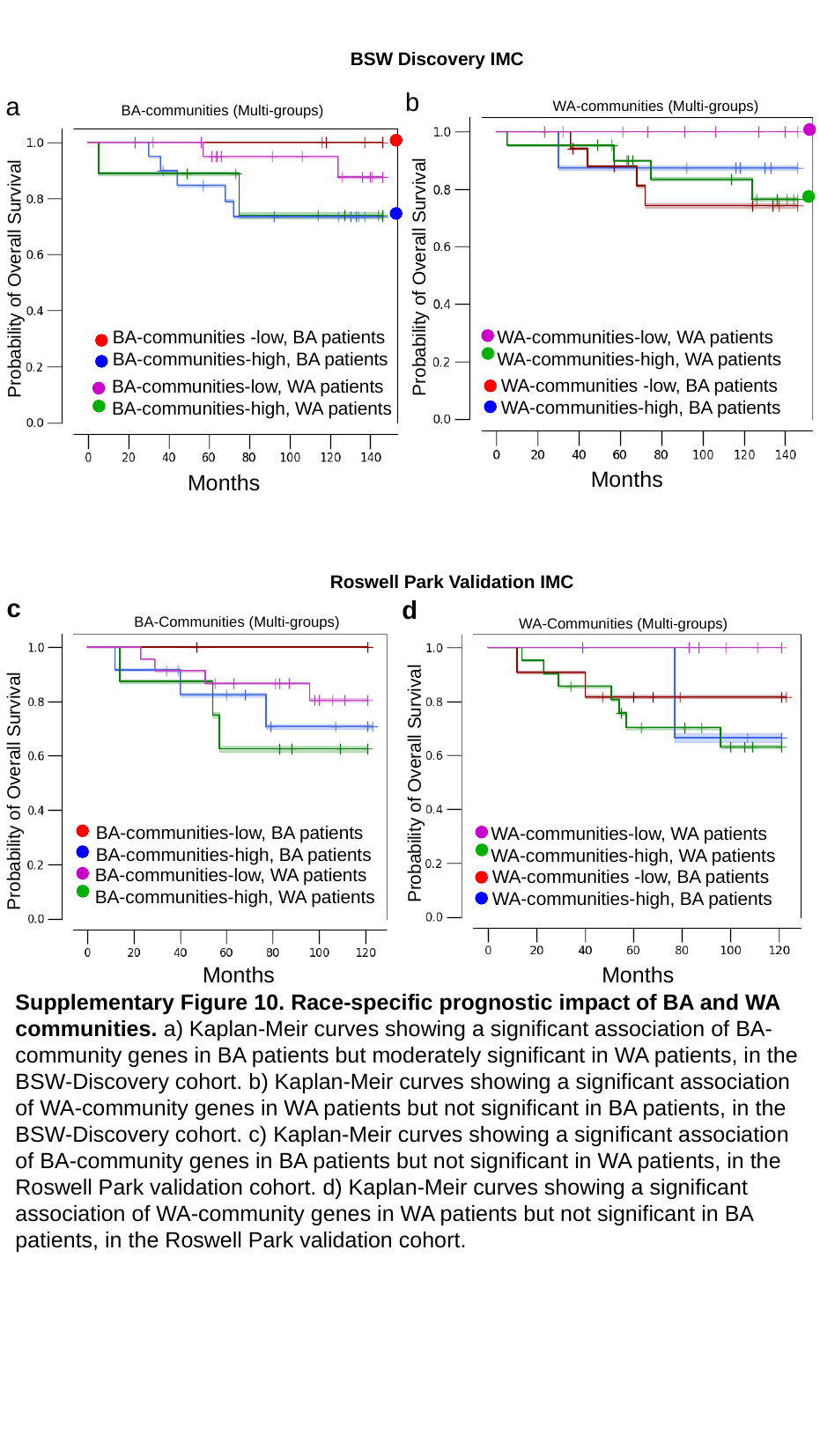

BSW Discovery IMC
b
a
WA-communities (Multi-groups)
WA-communities-low, WA patients
WA-communities-high, WA patients
BA-communities (Multi-groups)
BA-communities -low, BA patients
BA-communities-high, BA patients
Probability of Overall Survival
Probability of Overall Survival
BA-communities-low, WA patients
BA-communities-high, WA patients
WA-communities -low, BA patients
WA-communities-high, BA patients
Months
Months
Roswell Park Validation IMC
c
d
BA-Communities (Multi-groups)
WA-Communities (Multi-groups)
Probability of Overall Survival
Probability of Overall Survival
WA-communities-low, WA patients
WA-communities-high, WA patients
BA-communities-low, BA patients
BA-communities-high, BA patients
BA-communities-low, WA patients
BA-communities-high, WA patients
WA-communities -low, BA patients
WA-communities-high, BA patients
Months
Months
Supplementary Figure 10. Race-specific prognostic impact of BA and WA
communities. a) Kaplan-Meir curves showing a significant association of BA-community genes in BA patients but moderately significant in WA patients, in the BSW-Discovery cohort. b) Kaplan-Meir curves showing a significant association of WA-community genes in WA patients but not significant in BA patients, in the BSW-Discovery cohort. c) Kaplan-Meir curves showing a significant association of BA-community genes in BA patients but not significant in WA patients, in the Roswell Park validation cohort. d) Kaplan-Meir curves showing a significant association of WA-community genes in WA patients but not significant in BA patients, in the Roswell Park validation cohort.

### Slide 12
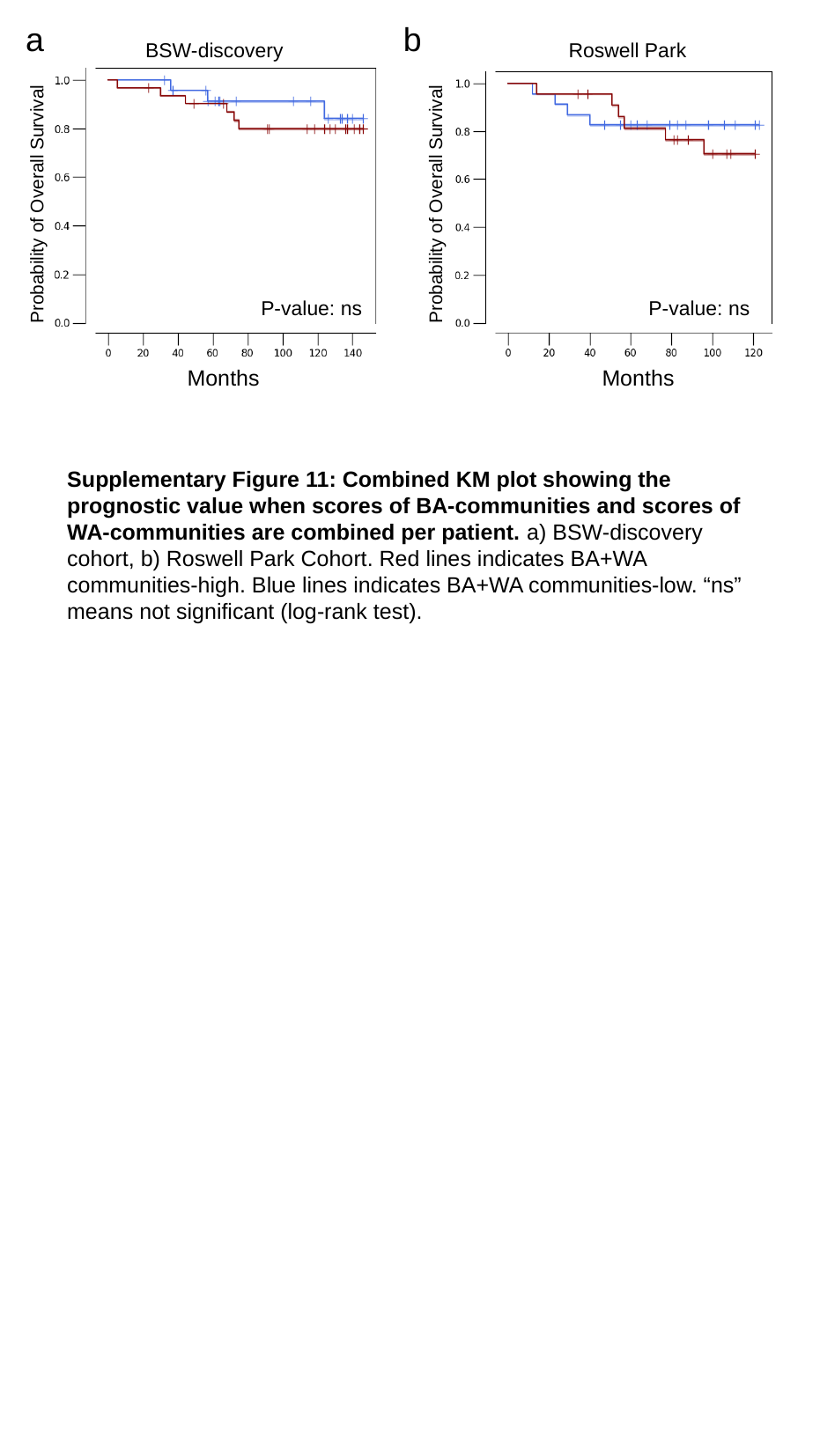

a b
BSW-discovery Roswell Park
Probability of Overall Survival
Probability of Overall Survival
P-value: ns
P-value: ns
Months
Months
Supplementary Figure 11: Combined KM plot showing the prognostic value when scores of BA-communities and scores of WA-communities are combined per patient. a) BSW-discovery cohort, b) Roswell Park Cohort. Red lines indicates BA+WA communities-high. Blue lines indicates BA+WA communities-low. “ns” means not significant (log-rank test).

### Slide 13
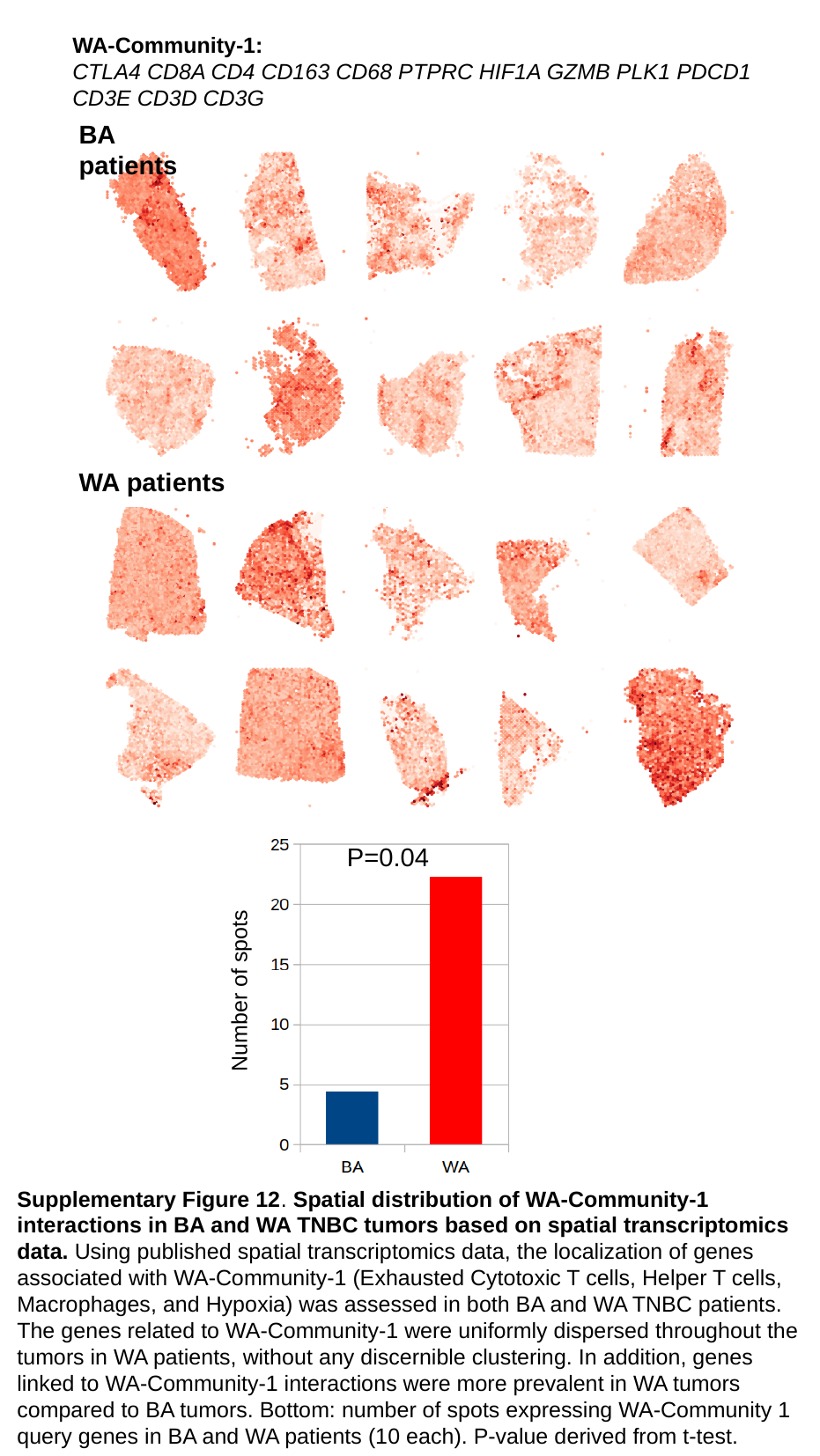

WA-Community-1:
CTLA4 CD8A CD4 CD163 CD68 PTPRC HIF1A GZMB PLK1 PDCD1 CD3E CD3D CD3G
BA patients
WA patients
P=0.04
Number of spots
Supplementary Figure 12. Spatial distribution of WA-Community-1
interactions in BA and WA TNBC tumors based on spatial transcriptomics
data. Using published spatial transcriptomics data, the localization of genes associated with WA-Community-1 (Exhausted Cytotoxic T cells, Helper T cells,
Macrophages, and Hypoxia) was assessed in both BA and WA TNBC patients.
The genes related to WA-Community-1 were uniformly dispersed throughout the
tumors in WA patients, without any discernible clustering. In addition, genes linked to WA-Community-1 interactions were more prevalent in WA tumors compared to BA tumors. Bottom: number of spots expressing WA-Community 1 query genes in BA and WA patients (10 each). P-value derived from t-test.

### Slide 14
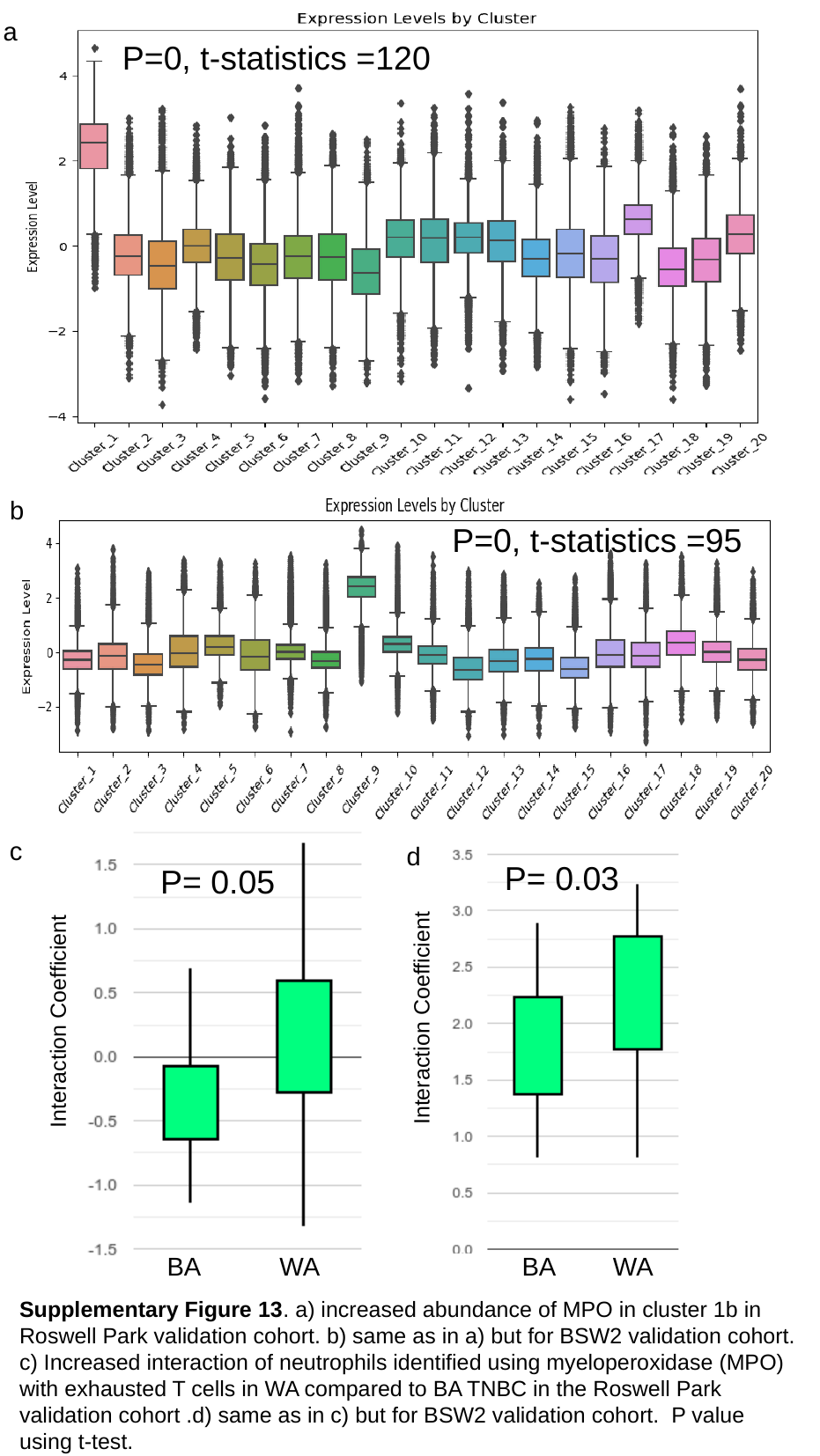

a
P=0, t-statistics =120
b
c
P=0, t-statistics =95
c
d
P= 0.03
P= 0.05
Interaction Coefficient
Interaction Coefficient
BA WA BA WA
Supplementary Figure 13. a) increased abundance of MPO in cluster 1b in Roswell Park validation cohort. b) same as in a) but for BSW2 validation cohort. c) Increased interaction of neutrophils identified using myeloperoxidase (MPO) with exhausted T cells in WA compared to BA TNBC in the Roswell Park validation cohort .d) same as in c) but for BSW2 validation cohort. P value using t-test.

### Slide 15
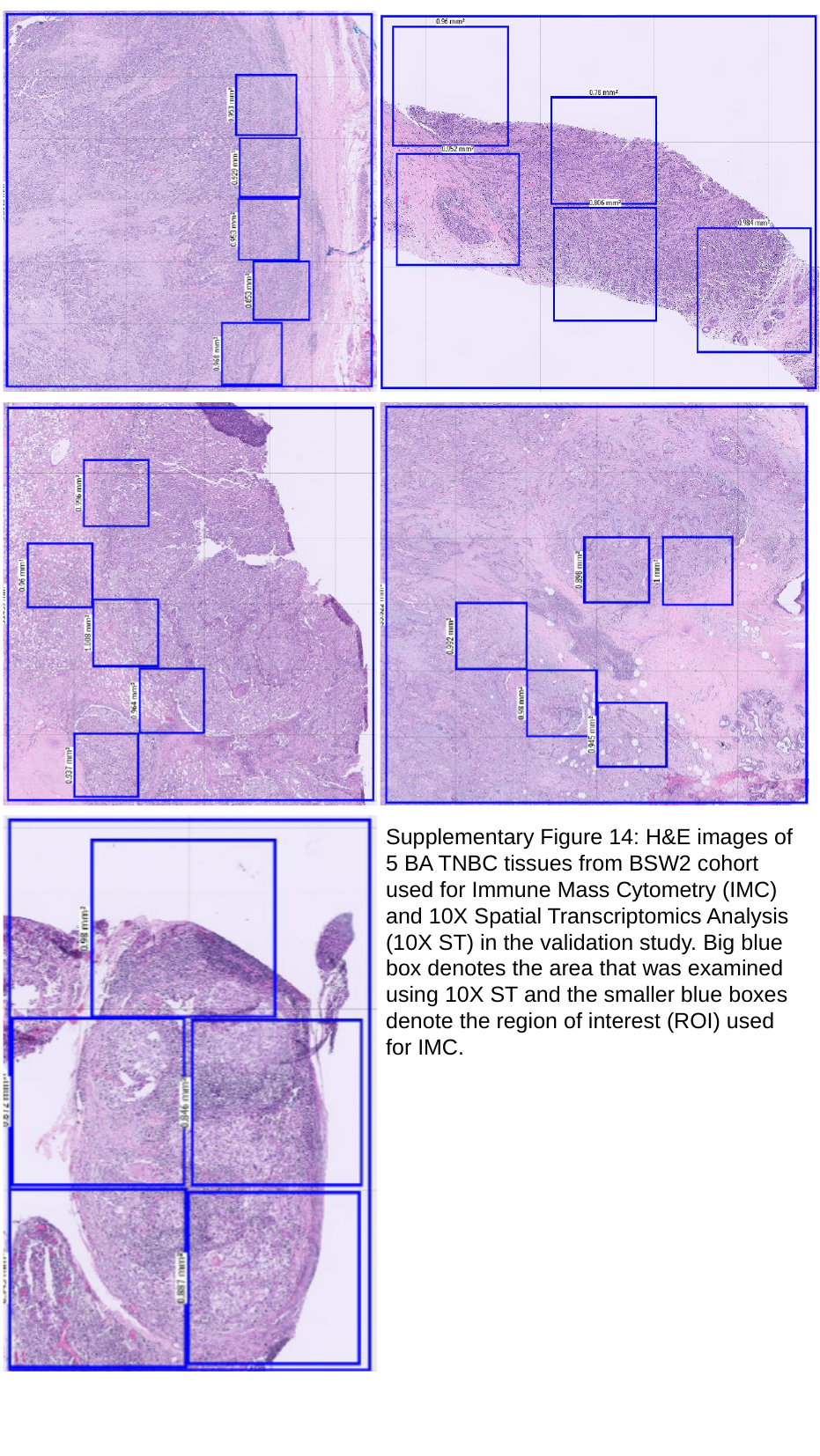

Supplementary Figure 14: H&E images of 5 BA TNBC tissues from BSW2 cohort used for Immune Mass Cytometry (IMC) and 10X Spatial Transcriptomics Analysis (10X ST) in the validation study. Big blue box denotes the area that was examined using 10X ST and the smaller blue boxes denote the region of interest (ROI) used for IMC.

### Slide 16
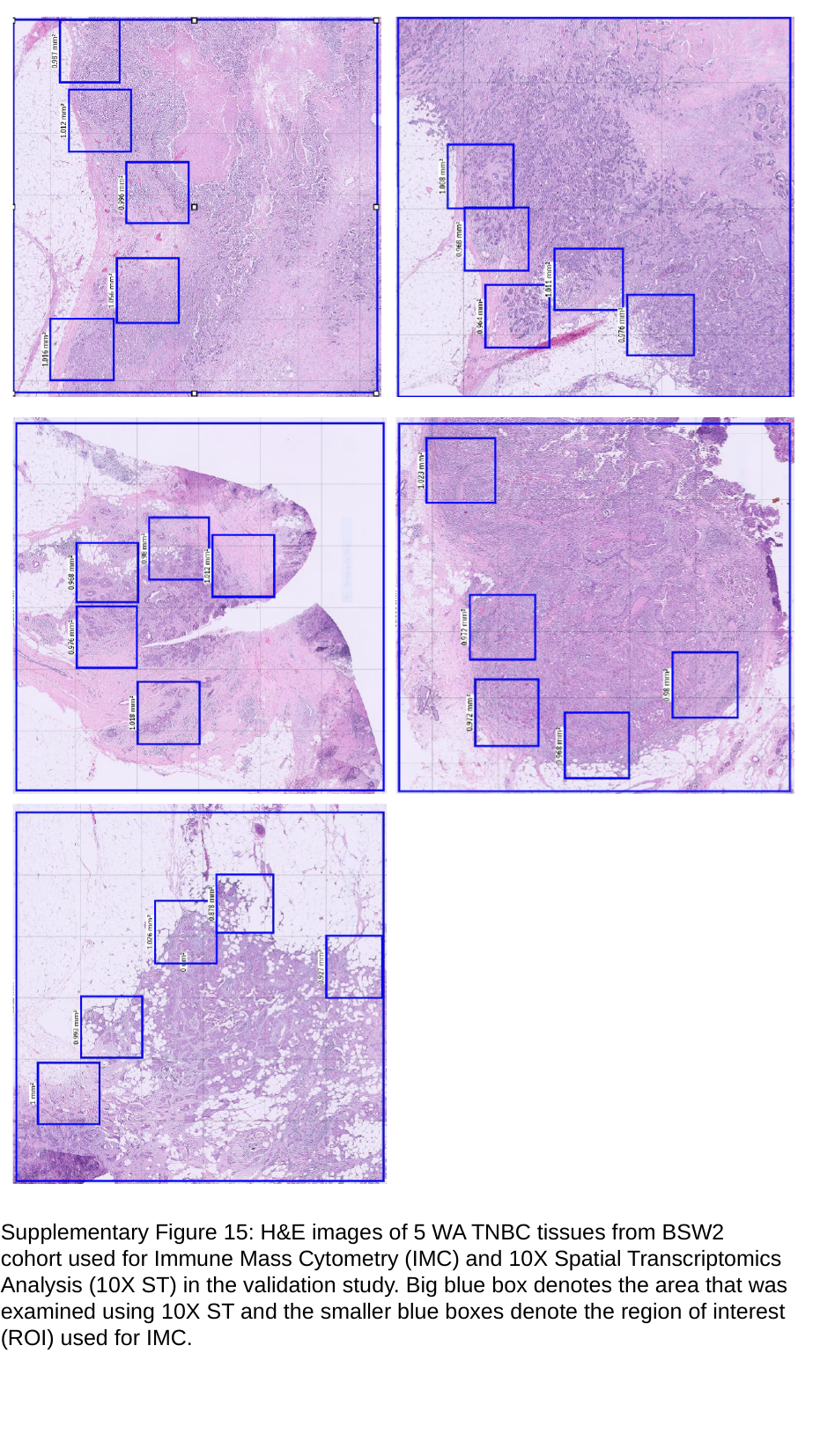

Supplementary Figure 15: H&E images of 5 WA TNBC tissues from BSW2 cohort used for Immune Mass Cytometry (IMC) and 10X Spatial Transcriptomics Analysis (10X ST) in the validation study. Big blue box denotes the area that was examined using 10X ST and the smaller blue boxes denote the region of interest (ROI) used for IMC.

### Slide 17
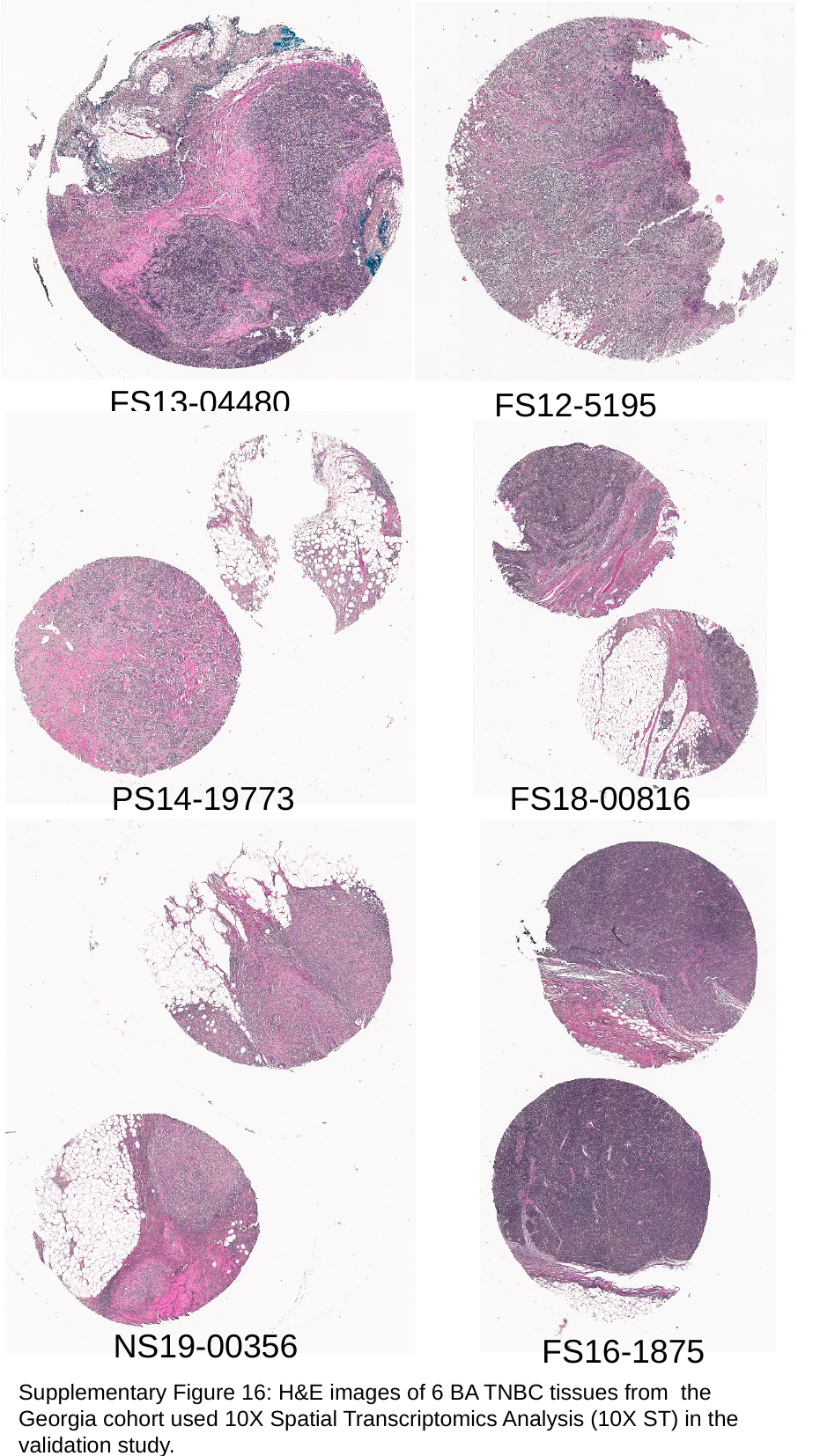

FS13-04480
FS12-5195
PS14-19773
FS18-00816
NS19-00356
FS16-1875
Supplementary Figure 16: H&E images of 6 BA TNBC tissues from the Georgia cohort used 10X Spatial Transcriptomics Analysis (10X ST) in the validation study.

### Slide 18
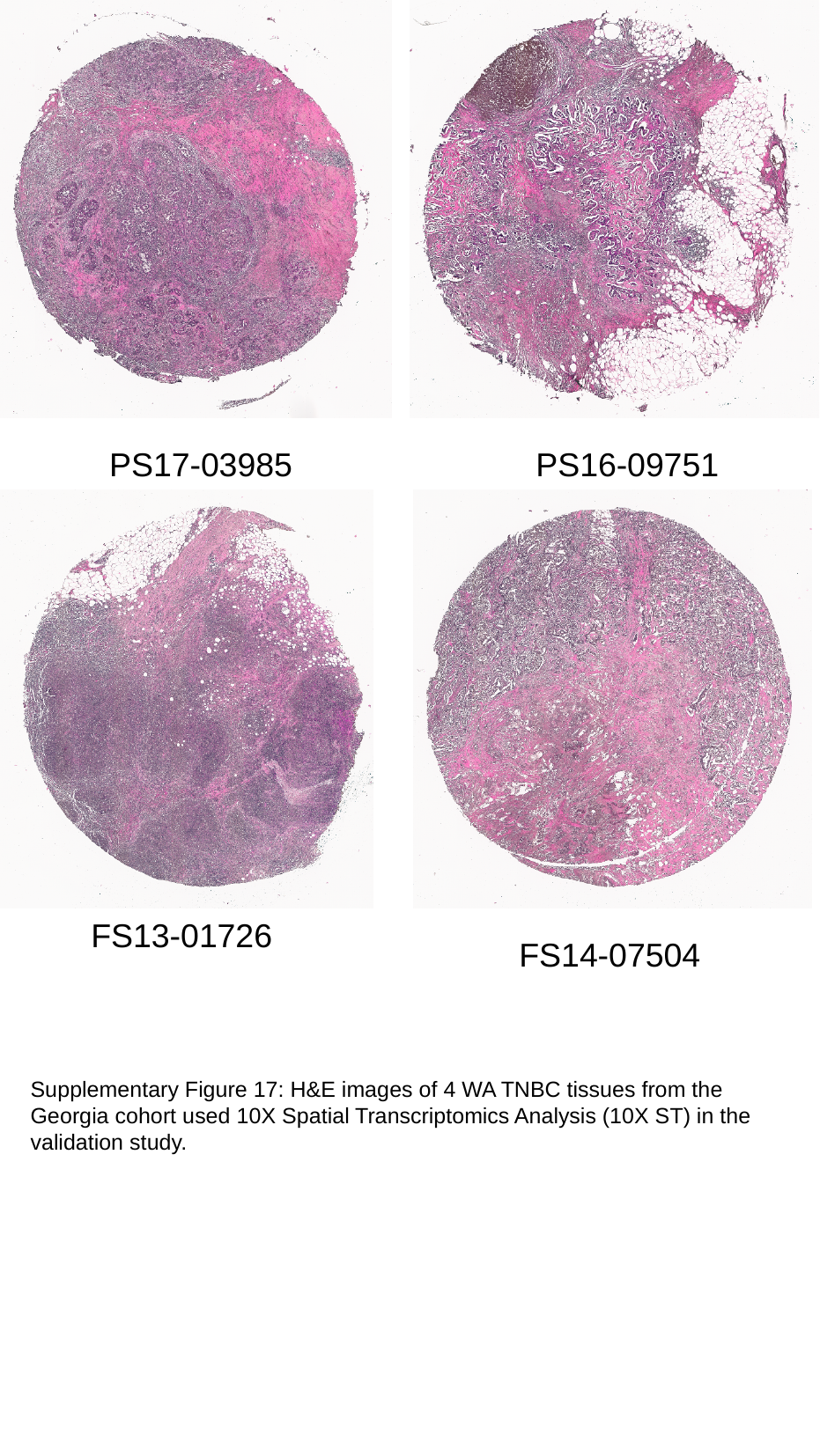

PS17-03985
PS16-09751
FS13-01726
FS14-07504
Supplementary Figure 17: H&E images of 4 WA TNBC tissues from the Georgia cohort used 10X Spatial Transcriptomics Analysis (10X ST) in the validation study.

### Slide 19
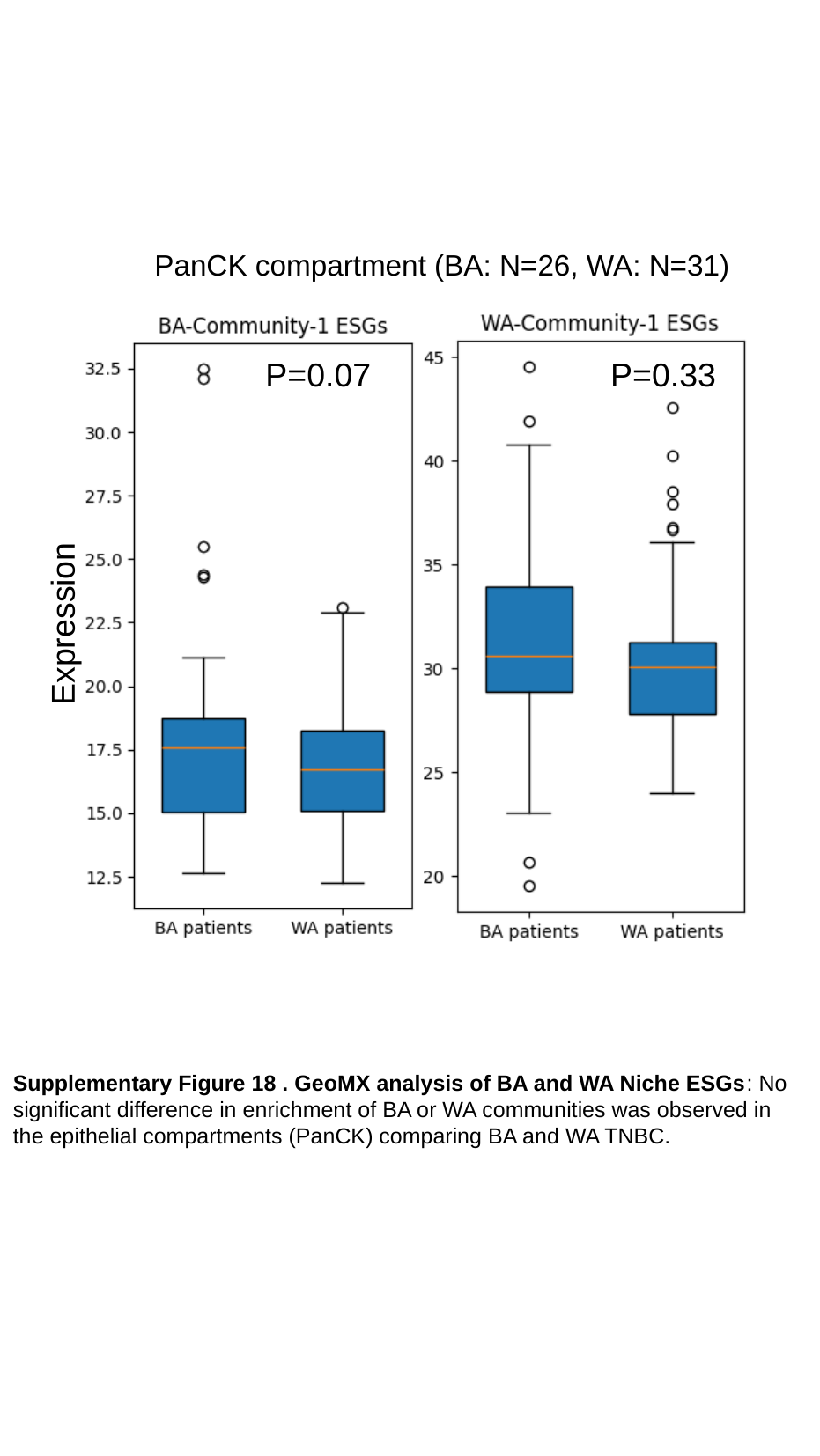

PanCK compartment (BA: N=26, WA: N=31)
Expression
P=0.07 P=0.33
Supplementary Figure 18 . GeoMX analysis of BA and WA Niche ESGs: No significant difference in enrichment of BA or WA communities was observed in the epithelial compartments (PanCK) comparing BA and WA TNBC.

### Slide 20
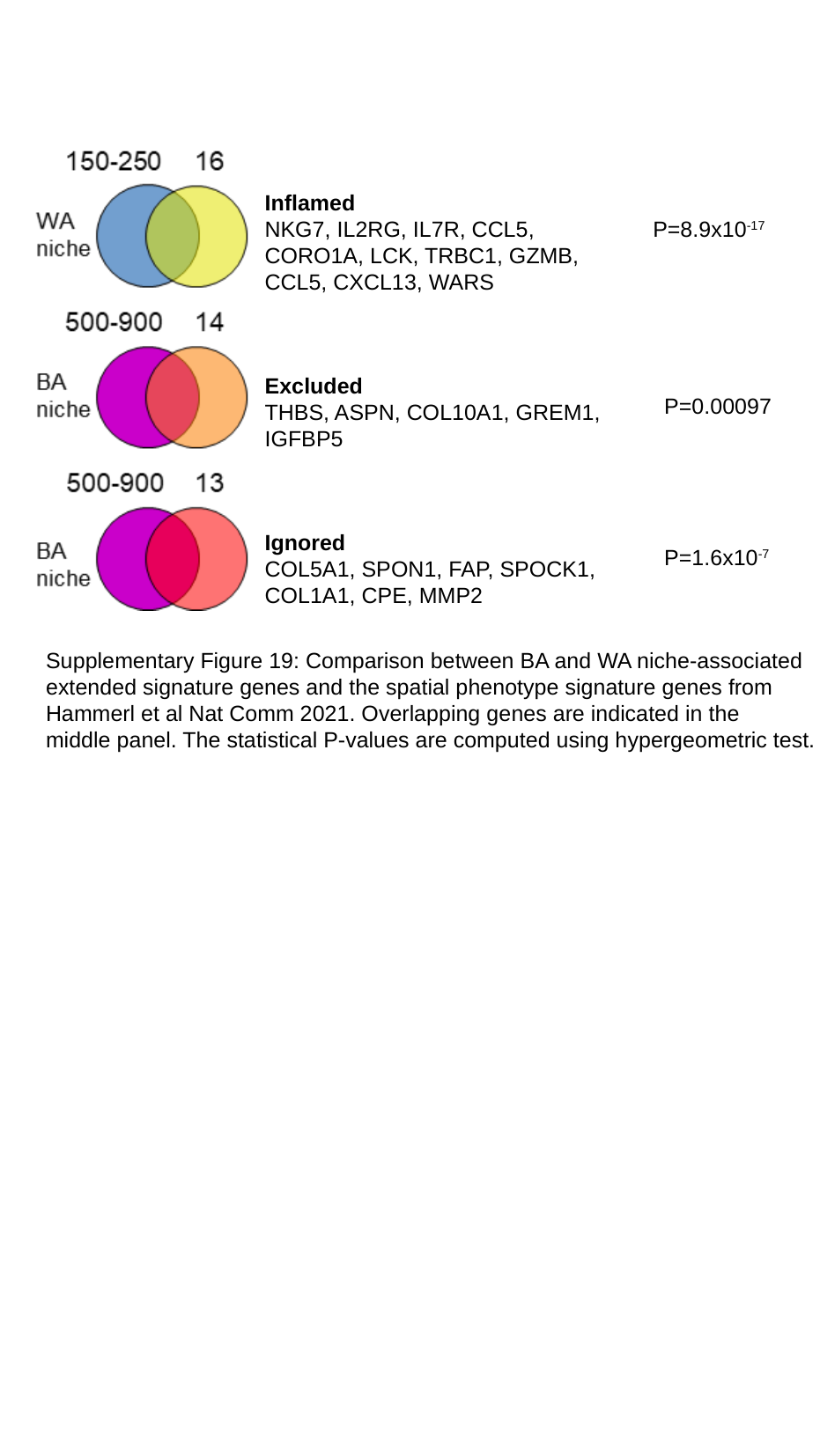

Inflamed
NKG7, IL2RG, IL7R, CCL5, CORO1A, LCK, TRBC1, GZMB, CCL5, CXCL13, WARS
P=8.9x10^-17^
Excluded
THBS, ASPN, COL10A1, GREM1, IGFBP5
P=0.00097
Ignored
COL5A1, SPON1, FAP, SPOCK1, COL1A1, CPE, MMP2
P=1.6x10^-7^
Supplementary Figure 19: Comparison between BA and WA niche-associated
extended signature genes and the spatial phenotype signature genes from
Hammerl et al Nat Comm 2021. Overlapping genes are indicated in the
middle panel. The statistical P-values are computed using hypergeometric test.
